# Feather keratin in *Pavo cristatus*: A tentative structure

**DOI:** 10.1101/2024.09.08.611866

**Authors:** Peter Russ, Helmut O. K. Kirchner, Herwig Peterlik, Ingrid M. Weiss

**Affiliations:** IBBS, University of Stuttgart, Pfaffenwaldring 57, D-70569 Stuttgart, Germany; Faculty of Physics, University of Vienna, Boltzmanngasse 5, A-1090 Vienna, Austria; SRF AMICA, University of Stuttgart, Pfaffenwaldring 32, 70569 Stuttgart, Germany; Suttgart Research Center Systems Biology (SRCSB), University of Stuttgart, Germany

## Abstract

The filament of the F-keratin polymer is an alternating arrangement of two tetrameric sequence segments, the “N-block” made of four strands AA 1–52, a twisted parallelepiped and the “C-block”, a sandwich of four strands AA 81–100. The N-blocks have 89°internal rotation within eight levels of *β*-sandwiches strengthened by three disulfide bonds per monomer. The C-blocks contain 5 aromatic residues, they provide resilience, like vertebral discs in a spinal column. The pitch of an N+C-block octamer is 10 nm. Solidification of F-keratin may involve the “C-blocks” to temporarily mold into “C-wedges” of 18° tilt, which align the polymer filaments into laterally amorphous fiber-reinforced composites of 9.5 nm axial periodicity. This distance corresponds to the length of the fully stretched AA 53–80 matrix segment. The “spinal column” is deformed like in scoliosis and unwinds under compression when F-keratin filaments perfectly align horizontally and form stacked sheets in the solid state.

## 1 Introduction

### 1.1 F-Keratin

F-keratin is a prerequisite for allowing birds to fly [1]. It forms composites with excellent mechanical, optical, and thermal properties. The F-keratin protein consists of 100 amino acids (AA) with many highly conserved regions among all birds. Evolutionary, F-keratin is a late invention, dating from 220-125 million years ago, when skin and integumentae of reptiles transformed into feathers [2, 3]. The peacock *Pavo cristatus* grows a 1 m long feather in about 100 days, at a rate of 100 nm/s [4].

### 1.2 AA sequence of the monomer

The 100 AA sequence (10,04 kDa) from *Pavo cristatus* F-keratin is shown in Figure 1 in a suggestive configuration, according to AA composition and hypothetical functions in three segments: AA 1–52 (N-block), AA 53–80 (GSA-string), and AA 81–100 (C-block). The most prominent feature is the presence of 9 cysteines, six of which are located in the N-block and 3 are located in the C-block. The C-block contains 25% aromatic residues. Besides the peacock’s (*Pavo cristatus*) F-keratin, we also investigated silver gull (*Chroicocephalus novaehollandiae*, also known as *Larus novaehollandiae*), and chicken (*Gallus gallus*) F-keratin. The highest degree of variability between species is in the GSA-string between AA 53–78 (Figure 2). It features mostly small and flexible residues (Gly, Ser, Ala), of which 8/27 = 30% are serins, similar to silk-like proteins. There is also considerable variation near the C-terminus, AA 79–100. In the peacock, it contains five aromates and three cysteines. Residues with key functions (Figure 2) are conserved in the same segments in all species, as is the length of the GSA-string, which fully stretched, would be 9.5 nm of the 35 nm length for the fully stretched whole sequence.

**Figure 1:**
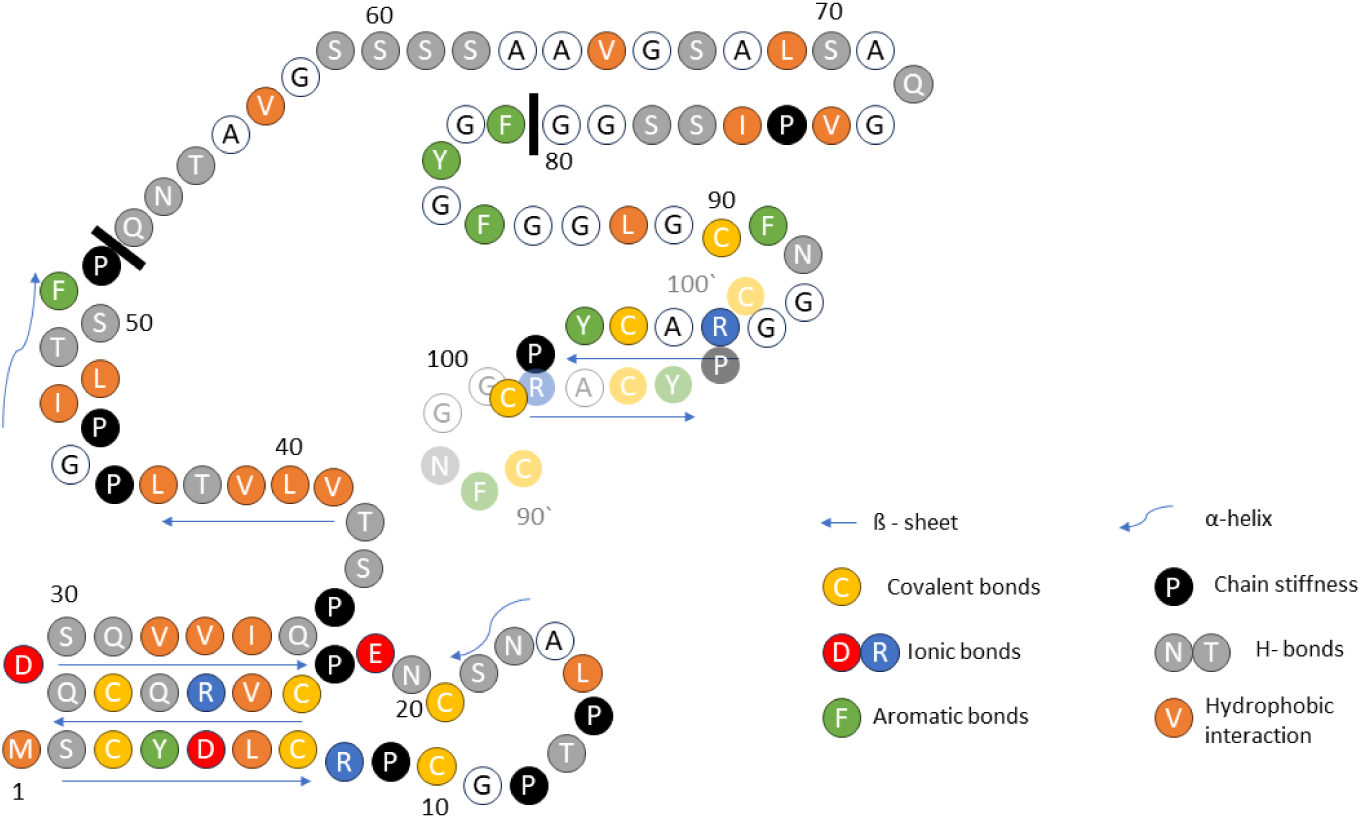
AA sequence of F-keratin from *Pavo cristatus* (UniProt accession number A0A8C9L9X5). Structural features with relevance to the new model are highlighted in color. The three functional parts of the primary structure are separated by solid black bars: N-block AA 1–52; GSA-string AA 53–80; and C-block AA 81–100. At the C-terminus, AA 90–100, the proximity of the Cys-90 to Cys-100 of another monomer (AA 90‘–100‘, transparent) is suggested by the disulfide-stabilized intermolecular *β*-sheet. Arrows indicate intra- and intermolecular *β*-sheets.

**Figure 2:**
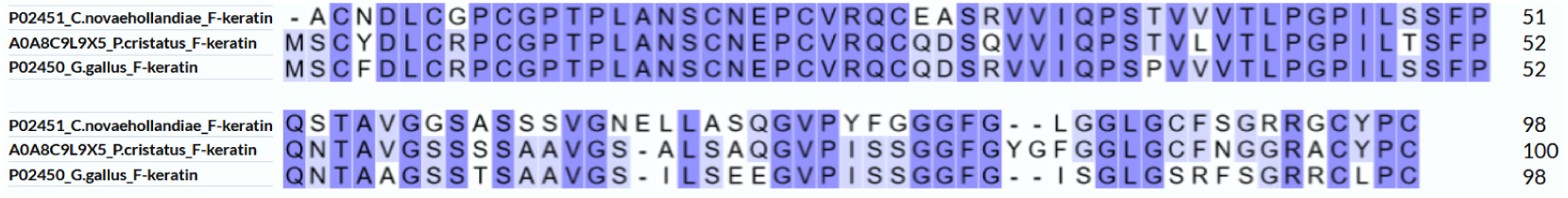
Sequences of silver gull (*Larus novaehollandiae*), peacock (*Pavo cristatus*), and chicken (*Gallus gallus*) (see Supplementary Materials 5.1 for UniProt accession numbers). The N-blocks AA 1–52, rich in Cys, are almost identical in the three species. The GSA-strings AA 53–80, rich in Gly, are randomly different. The C-blocks, AA 81–100, are dominated by aromates in all three.

### 1.3 Diffraction data

The molecular structure of the rachis of the peacock‘s tail feather is highly ordered, as obvious from the diffraction patterns shown in Figure 3 [4]. However, no definite crystal structure of F-keratin could be derived. The X-ray pattern in different parts of the rachis is highly stable, it changes only slightly across the thickness [5] and along the length of 1 m from the calamus to the tip, both in *Pavo cristatus* and *Pavo cristatus mut. alba* [4].

**Figure 3:**
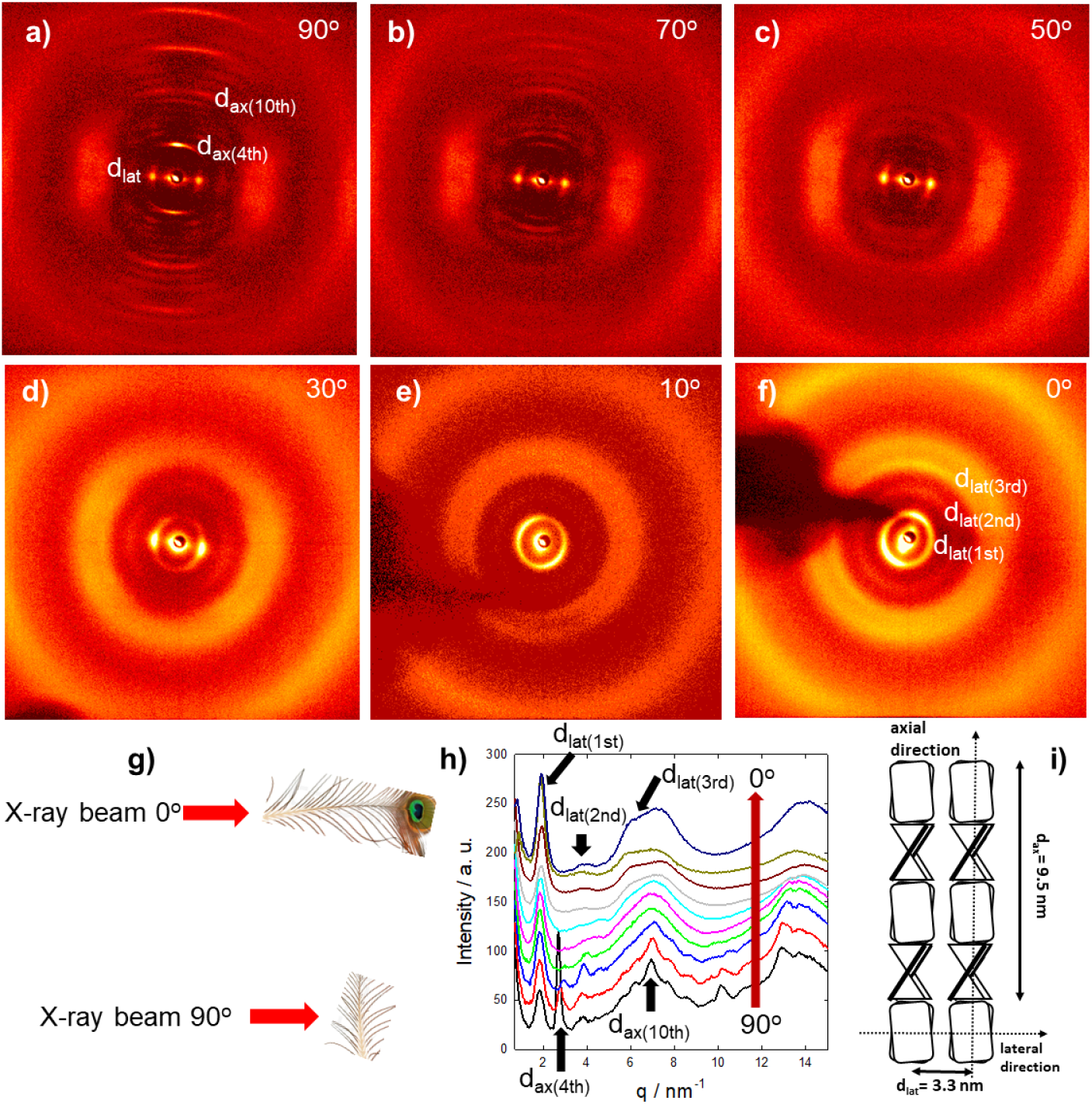
Diffraction patterns taken at different rotation angles: (a-f) X-ray beam perpendicular to the fiber axis, (a) 90°, to X-ray beam in fiber axis, (f) 0°. (g) Measurement geometry. (h) Integrated diffraction intensities at different angles (red arrow, 90°to 0°). (i) Model adapted from [4]. The main axial reflections, arising from the meridional repeat distance of 9.4 nm, disappear from 90° to 0°(a-f), whereas the main lateral spot from the lateral repeat distance of 3.4 nm develops into a ring. Also, higher lateral reflections appear (e, f). Distances in real space are d = 2*π/q*, the first peak at 1.85 nm*^−^*^1^ corresponds to 3.4 nm, the second peak at 2.7 nm*^−^*^1^ to 2.35 nm.

Selected X-ray patterns are shown in Figure 3a-f, taken at different X-ray beam angles from 90° (= perpendicular to the rachis axis) to 0° (= aligned in the rachis axis) as indicated in Figure 3g. The integrated intensities for all rotation angles measured in ten degree steps are presented in Figure 3h. The 90°configuration measurement shows a highly ordered structure with lots of diffraction peaks arising from the axial arrangement of F-keratin with a repeat distance of 9.4 nm. The strong diffraction spot from the 4th layer line at 9.4/4 *≈* 2.35 nm vanishes already in the 70°configuration. At 40°and less, only the lateral peaks are visible. The lateral spot at the 90° configuration develops into a ring at the 0°configuration. Thus, there is no preferred orientation of the filaments in the plane perpendicular to the long axis of the feather. Three further rings could be identified, arising from a repeat distance of 3.4 nm. Rotating the rachis with respect to the X-ray beam clearly shows the long-range order of the molecules within the F-keratin filaments along the feather axis and a weak order in the plane normal to it. The structural model for peacock F-keratin in Figure 3i [4] is based on the pioneering work on seagull F-keratin [6–9] and claims of Fraser and Parry [10–19]. The axial assembly of the previously proposed *β*-sandwiches arises from a four-fold screw axis, which generates a filament with a pitch length P in the seagull of about 9.5 nm [8] and an axial rise of about 2.4 nm [6]. If the filaments are densely packed, the diameter and the distance between filaments in the lateral direction coincides and was measured to be 3.4 nm [9], also in the rachis of the peackock‘s feather [4]. Small changes in P have been observed in some keratinous tissues (9.2 nm in chicken scale; 9.28 nm in snake scale; 9.85 nm in lizard claw [12]), but the four-fold screw symmetry over P is maintained. It would seem probable, therefore, that the basic helical framework of the filaments is conserved across all of the sauropsids [13].

### 1.4 AlphaFold

Since 2021, the AlphaFold algorithm has been available for predicting protein folding from the amino acid sequence, the so-called primary structure [20]. This algorithm can even predict unknown 3D model structures for proteins with low sequence similarity to deposited experimental structures in the PDB database [20]. By its very nature, AlphaFold has a preference for structures similar to those already in its library, it seeks familiar and avoids unfamiliar structures. It is eminently suitable for functional proteins, less so for structural proteins. The reason is that monomers of a structural protein are not functional as individual monomers in aqueous solvents, structural proteins agglomerate or crystallize. F-keratin is such an example of a structural protein that likely transforms into polymeric filaments or fibrils and even further into solids. AlphaFold cannot handle collective agglomeration, but can furnish valuable information and insight into parts of such structural entities, when essentially assisted by guesswork, chemical, biological, and physical intuition. None of the F-keratin structures, listed in the database under www.alphafold.ebi.ac.uk fit known diffraction data. Our goal was to take advantage of the AlphaFold algorithm for predicting a reasonable structure, which fits experimental structural and mechanical data of *Pavo cristatus* (Supplemenatry Materials 5.1 and 5.2). Here we report the novel structure, only parts of which were furnished by AlphaFold, the rest being derived from X-ray, chemical, and mechanical data, as well as the usage of further bioinformatic tools like ClusPro and GROMACS.

## 2 Results

### 2.1 Functional regions

AlphaFold, when asked to arrange full-length sequences AA 1–100 as a monomer, dimer, tetramer, or octamer did not produce any reasonable structure, but it does so for parts of the sequence. In trials with segments of various lengths and positions, only a few results passed visual inspection, where the shapes could be stacked together in a meaningful way. Remarkably, these reflect the compositional inhomogeneity in three parts of the AA 1–100 sequence, as shown in Figure 1. AlphaFold produced a convincing result, with the highly conserved AA 1–52 segment only as a tetramer (section 2.2, Figure 4). For the AA 81–100 segments, reasonable structures were obtained for dimers and tetramers (section 2.3, Figure 5). For the AA 53–80 segments, AlphaFold produced only variations of helices and random coils.

**Figure 4:**
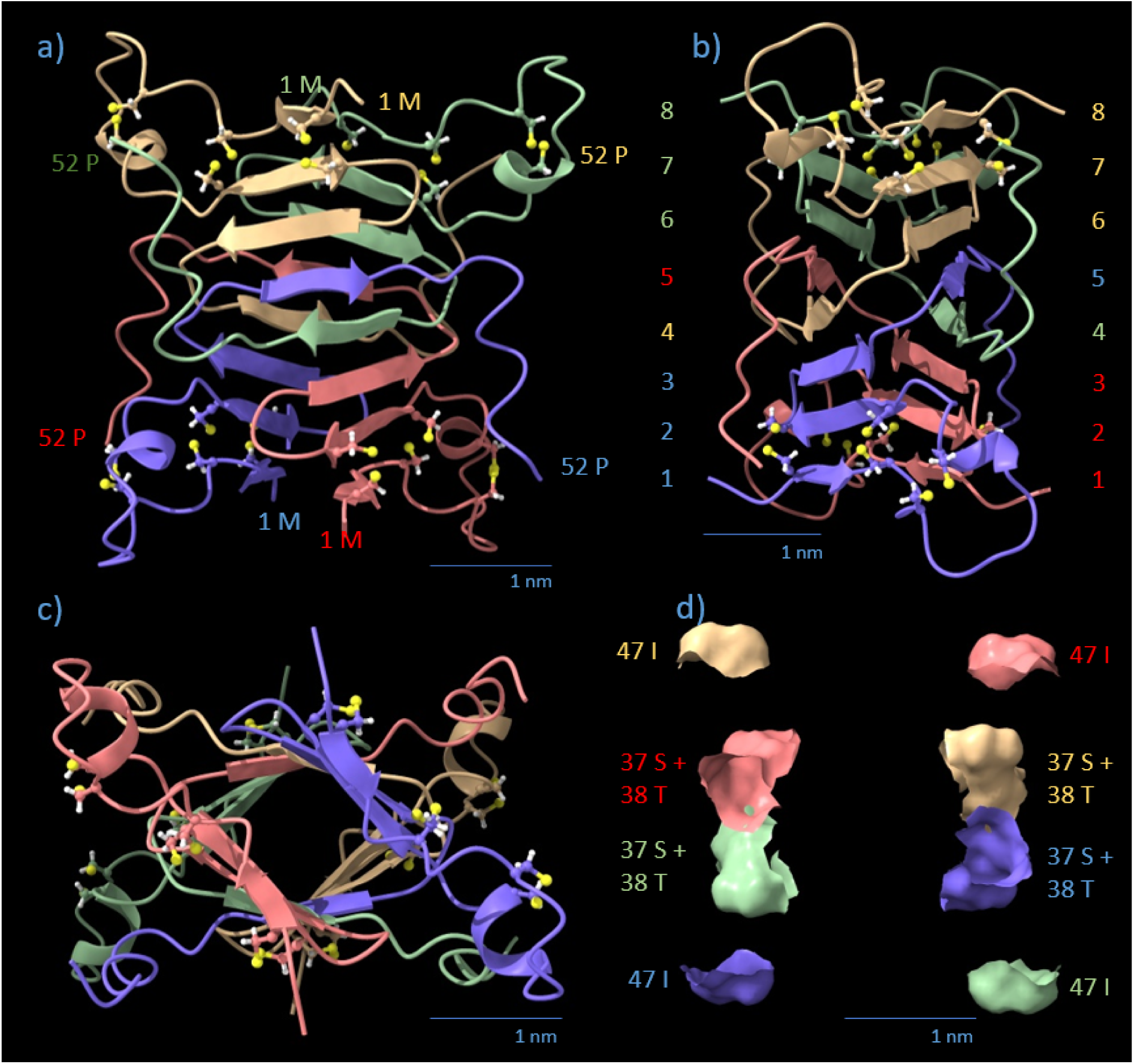
AlphaFold prediction of most parsimonious tentative structure of the F-keratin tetrameric N-block, AA 1-–52 (4 *×* 52 = 208 residues). Monomers are color-coded by chain (blue, red, green, yellow). Cysteine residues are displayed in a ball-stick view with standard colors. (a) Side view, staircase arrangement of helical *β*-strands in levels 1-8 (highlighted in the colors of adjacent *β*-strands) in the axial direction. Chain orientations of individual monomers are indicated by the respective AA number. (b) side view 90° turned, *β*-strands are rotationally staggered by an average horizontal angle of 11.125°per *β*-strand. The 8th strand is rotated 89°against the 1st strand in *Pavo cristatus*. The distance between the sandwiched sheets is 1.0-1.2 nm. Note that the polypeptide backbones of the four monomers are intertwined between strands in levels 4 and 5. (c) Axial view. (d) Equatorial cross-section of (a), corresponding to (c). AA 37-S, 38-T, and 47-I sit in the equatorial plane.

**Figure 5:**
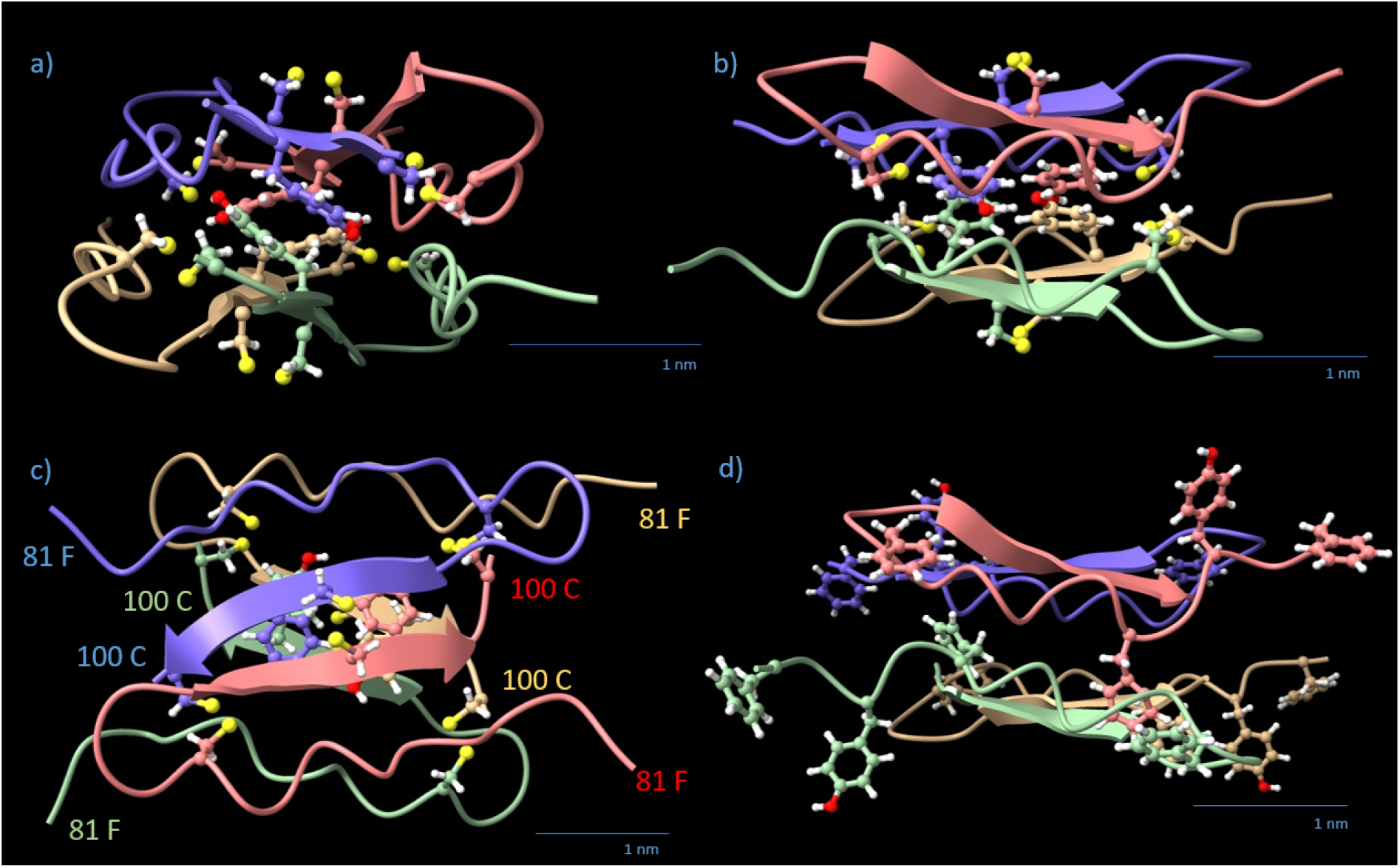
AlphaFold prediction of the *Pavo cristatus* F-keratin tetrameric C-block, AA 81-–100 (4 *×* 20 = 80 residues). Monomers are color-coded by chain (blue, red, green, yellow). The four 98-Y and 12 cysteine residues are displayed in a ball-stick view with standard colors. Chain orientations of individual monomers are indicated by the respective amino acid number. (a) Side view, filament axis vertical. (b) The side view turned 90°around the filament axis. (c) Top view, along the filament axis. Note that each pair of *β*-strands forms one level; the arrangement is rectangular rather than square in the (x,y) plane. The direction of the *β*-strand arrows is from N-to C-terminus. (d) shows the outer aromates from the 81-FGYGFGGLGCF motif in the same view as (b).

One promising tetrameric structure of the AA 1–52 segment formed two H-bond stabilized *β*-strands, covalently crosslinked by 3 intramolecular disulfide bonds between 6 Cys residues, a unique stiffening feature of mechanical importance. A second mechanical aspect concerns the AA 81–100 segment, which has the potential to form three intermolecular disulfide bonds (90– 100’, 97–97’, 100–90’). These stabilize an intermolecular *β*-sheet, between two AA 81–100 monomers. Two such dimers harbor 4 *×* 5 aromatic amino acids (3 F and 2 Y per monomer), which would influence the mechanical functionality by not being stiff but resilient, depending on the geometric arrangements of the side chains. One promising docking experiment suggested a vital role of 5-D and 95-R, which appear at the interface with a perfect steric fit, thus providing two strong ionic bonds between each tetrameric N-block and C-block. The AA 53–80 segment is rich in Gly, Ala, and Ser, which makes it soft in bending. Fully stretched, it could span 9.5 nm in length, similar to the repeating unit as identified by SAXS measurements. Based on all these observations, we decided to focus on this tentative molecular structure for F-keratin in the solid state.

### 2.2 N-block

The AlphaFold result for the N-block tetramer (4 *×* AA 1–52) is shown in Figure 4. The algorithm predicts four interpenetrating monomers in a twisted parallelepiped of the dimensions 3.0 nm *×* 3.9 nm *×* 4.0 nm (x,y,z). In z-orientation (filament axis), there are 8 levels occupied by 2 opposing antiparallel *β*-strands each. The levels are highlighted by numbers in Figure 4b in the respective colors of the monomer chains. The radial distance between opposing twisted *β*-sheets is 1.0-1.2 nm. The polypeptide strands cross two times each over the equatorial plane (z = 0) at the positions 37-S to 38-T within the helical *β*-core and 47-I in the outer *α*-helix (Figure 4d). Each monomer has many contact surfaces with any other monomer, with H-bonds, as well as hydrophobic interactions holding the core together (Supplementary Materials 5.6, Data S1, S2, and S3). The N-block transition regions AA 45–52 are folded into a flexibly linked *α*-helix which leaves the core horizontally at levels 4 and 5 of the helical *β*-core.

The AA 7–23, responsible for the external shape of the N-block, resemble a wing screw stabilized by 2 disulfide bridges per monomer and 4 proline residues. It defines the maximum diagonal dimension of the parallelepiped, spanning 4.3 nm in the xy-plane (Figure 4). Together with the transition *α*-helices (AA 45–52), the wing screws define the contact area for lateral spacing. The N-block displaces a volume V = 23.088 nm^3^ (57% of total volume), with a total mass of 22.32 kDa (107.3 Da/AA), resulting in a density of 0.96 kDa/nm^3^. The arrangement of the 8 levels of *β*-sheets in the central core of the N-block resembles, but is not, a left-handed helix (Figures 4a, 4b, and 4c). The upper half of the tetrameric N-block, 0 < z < 2.2 nm, twists 45°to the left. The lower half, - 2.2 nm < z < 0, twists another 45°to the right. The N-block is a monoclinic “twisted parallelepiped” (Figure 4d). The top and bottom surface planes are inclined concerning the z-axis. The twist is not strictly constant, but between z = - 2.2 nm and z = + 2.2 nm it is symmetric concerning z. It adds up to 89°(*±* 2.3°, measured in various projections) over the height of the N-block.

### 2.3 C-block

The most promising AlphaFold model of the C-terminal *Pavo cristatus* F-keratin segment AA 81-–100 is a flat rhombohedral tetrameric disc with dimensions 2.5 nm *×* 3.3 nm *×* 1.5 nm (x,y,z) (Figure 5). This C-block is slightly smaller than the N-block cross-section and fits sterically well between two of them. It contains 25% aromatic AA. The structure has four *β*-strands with the four 98-Y residues lined up in the central plane of the disc, forming two aromatic stacks [21–23]. The *β*-sheet core is surrounded by the 16 remaining aromatic residues 81-F, 83-Y, 85-F, and 91-F. For functional reasons, we divide the C-block into two regions: (1) the inner C-block hinge (key amino acid: 98-Y) and (2) the outer 81-FGYGFGGLGCF motif (”rubber cushion ring”). The disc displaces 7.935 nm^3^ (20% of total volume) with a mass of 8.2 kDa (102.5 Da/AA), resulting in a density of 1.033 kDa/nm^3^. The z-axis is not a mirror normal, but there is no distinction between the +z and -z directions. The .pdb file is included as Data S4 (Supplementary Materials 5.6).

### 2.4 GSA-String

The AA 53–80 segment does not yield any reproducible folding into secondary structures or 3D motifs when subjected to AlphaFold. The results were mostly elongated strings with partial random coil and *α*-helical elements. It is therefore called string. This region features mostly small and flexible AA residues such as glycine (Gly, G) or alanine (Ala, A). The 30% serine (Ser, S) fraction is as high as in spider silk [24]. It represents a common linker sequence [25]. The therefore called “GSA-strings” of 4 *×* 28 AA leave the N-block equatorial, forms a presumably amorphous polypeptide matrix and covalently connects the tetrameric N- and C-blocks within and/or between filaments. The 112 AA/tetramer displace a volume of 9.612 nm^3^ (24% of total volume) with a mass of 9.8 kDa (87.5 Da/AA) and a calculated density of 1.02 kDa/nm^3^.

### 2.5 The Monomers

The folded monomers were extracted from the folded *Pavo cristatus* F-keratin tetramer and superimposed. The aligned 3D structures of the 4 monomers are almost identical, taking computational inefficiencies into account (Figure 6a). 3D structure alignments of folded F-keratin N-block monomers built from *P. cristatus*, *L. novaehollandiae*, and *G. gallus* show the same principle structure (Figure 6b). Since each monomer structure has been individually reproduced, the observed similarity, even across species is a strong indication of the reliability of the AlphaFold algorithm in this particular case.

**Figure 6:**
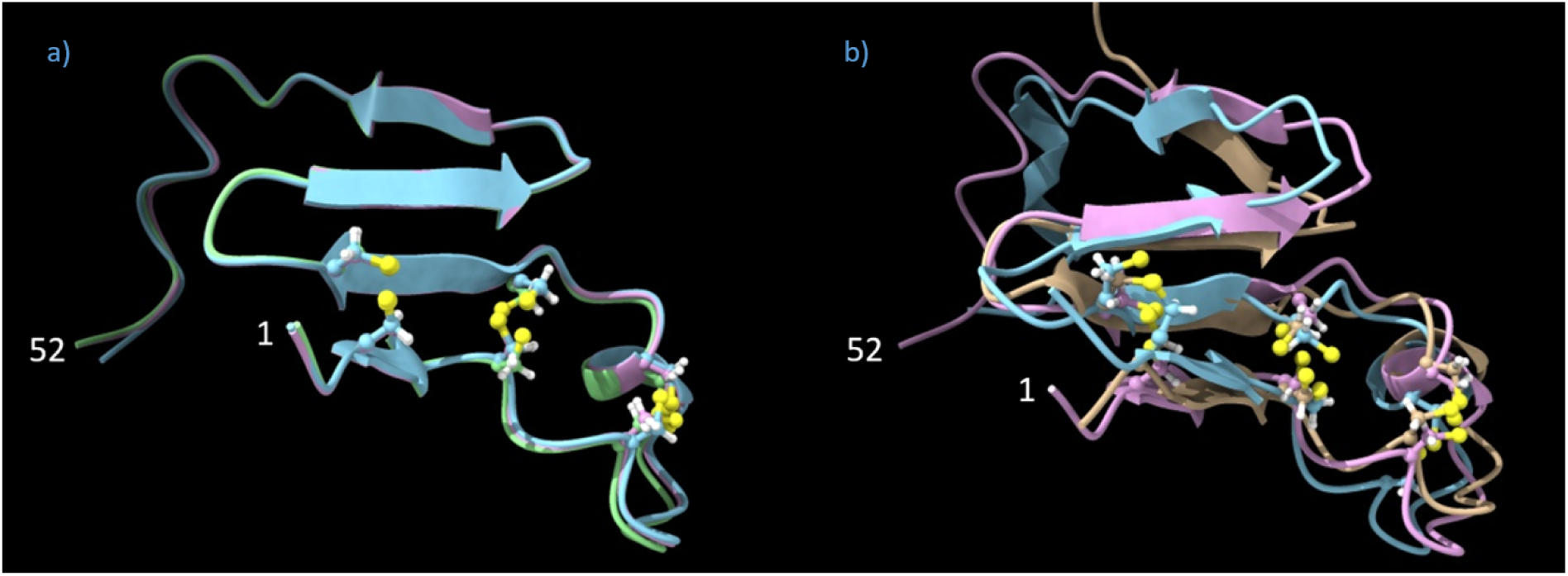
Structural alignments of extracted F-keratin monomer fragments, analyzed with jFATCAT. a: Overlay of three *Pavo cristatus* F-keratin N-block monomers (AA 1–52). The four *β*-sheets and the sulfur atoms (in yellow, ball-stick display) which link the cysteines at positions 3/27, 7/23, and 10/19 are perfectly aligned. b: Overlay of extracted F-keratin N-block (AA 1–52) chains from *P. cristatus*, *G. gallus*, and *L. novaehollandiae*.

Structural alignments of AlphaFold folded monomers from the C-block segment also produced reasonable results, with the exception that the characteristic shape of the disc, which perfectly fits the N-block interfaces for positively and negatively charged amino acid residues, was only obtained for the F-keratin sequence of *Pavo cristatus*. So far, according to AlphaFold models, C-block segments of *Gallus gallus* and *Larus novaehollandiae* lack such interfacial fit.

### 2.6 Assembly of the filament by stacking in z-direction

The *Pavo cristatus* F-keratin N-block and C-block are suitable for a regular arrangement into filaments, similar to the polymerization of actin [26]. The internal helical structure of the N-block causes a relative rotation of 89° *±* 2.3° between the bottom and top-level *β*-strands of each tetramer. The C-tetramer has no internal rotation but has a fixed orientation relative to the N-blocks on both sides due to two pairs of specific ionic bonds (5-D/95-R). The GSA-string does not contribute to any specific structure. The polymer stacking (0°-N-block*_n_* - 89°/ C-block / 89°-N-block*_n_*_+1_ - 178°/ C-block / 178°-N-block*_n_*_+2_ - 267°/ C-block / 267°-N-block*_n_*_+3_ - 356°) is the self-evident consequence. Over the period of two N-block/C-block tetramers, it provides *≈* 180° (178°) rotation, which is equivalent to *≈* 360°(356°) rotation because of the D2 symmetry [27]. The period of the resulting octamer is 10 nm. The closest match with experimental data would be the 9.4 nm SAXS signal. The 8 GSA-strings AA 53–80 have no assigned role, they exit the tetrameric N-blocks at monomer position AA 53 and enter the tetrameric C-blocks at monomer position AA 80. If fully stretched, the GSA-string might connect vertically to another octamer unit of the same F-keratin filament, or cross-link horizontally to a core unit of neighboring F-keratin filaments. Roughly estimated, the maximum reach of the GSA-string is about 9.5 nm.

The interfaces between tetrameric N-blocks and C-blocks are identical at all levels (Supplementary Materials 5.6, Data S5). In z-direction, there are 20 *β*-strand levels in the octameric N-block/C-block unit, 12 of which are clamped together by disulfide bridges (N-block levels 1,2,7,8, plus C-block levels 1 and 2; Figure 4, Figure 5). As verified by docking experiments, the C-block fits into the recess on both sides of the N-block, they perfectly connect the two elements via two electrostatic bonds from 5-D to 95-R* and 5-D’ to 95-R’* (*: any R-95 from monomers within 9.5 nm distance) on both sides. This is a remarkably perfect fit and excludes any other possibility of turning a C-block with respect to its neighboring N-block. If N-block and C-block are turned by 0° to *≈* 90°(89°) against each other, this turn must occur in the plane of the aromatic residues that connect the two halves of the disc. The total height of the repeating unit was 5.2 nm when manually measured from the aromatic rings of 98-Y of one C-block disc to the next, or 4.9 nm when measured from C*_α_* to C*_α_* (Figure 7). Unfortunately, it was not possible to simulate, by employing the docking strategy, a vertical “Velcro”-like intercalation of the four outer aromatic residues, the 81-FGYGFGGLGCF motif. Due to the steric flexibility, these aromatic residues may be squeezed out laterally from the main filament under compression. Due to the relatively large size of Tyr and Phe (main axes: Y, 1.04 nm; F, 0.97 nm), their orientation with respect to the x,y-plane as well as intercalation matters. In principle, an intercalation would suffice to explain the deviation of 10 nm (in silico; C*_α_* to C*_α_*) - 9.4 nm (SAXS) = 0.6 nm.

**Figure 7:**
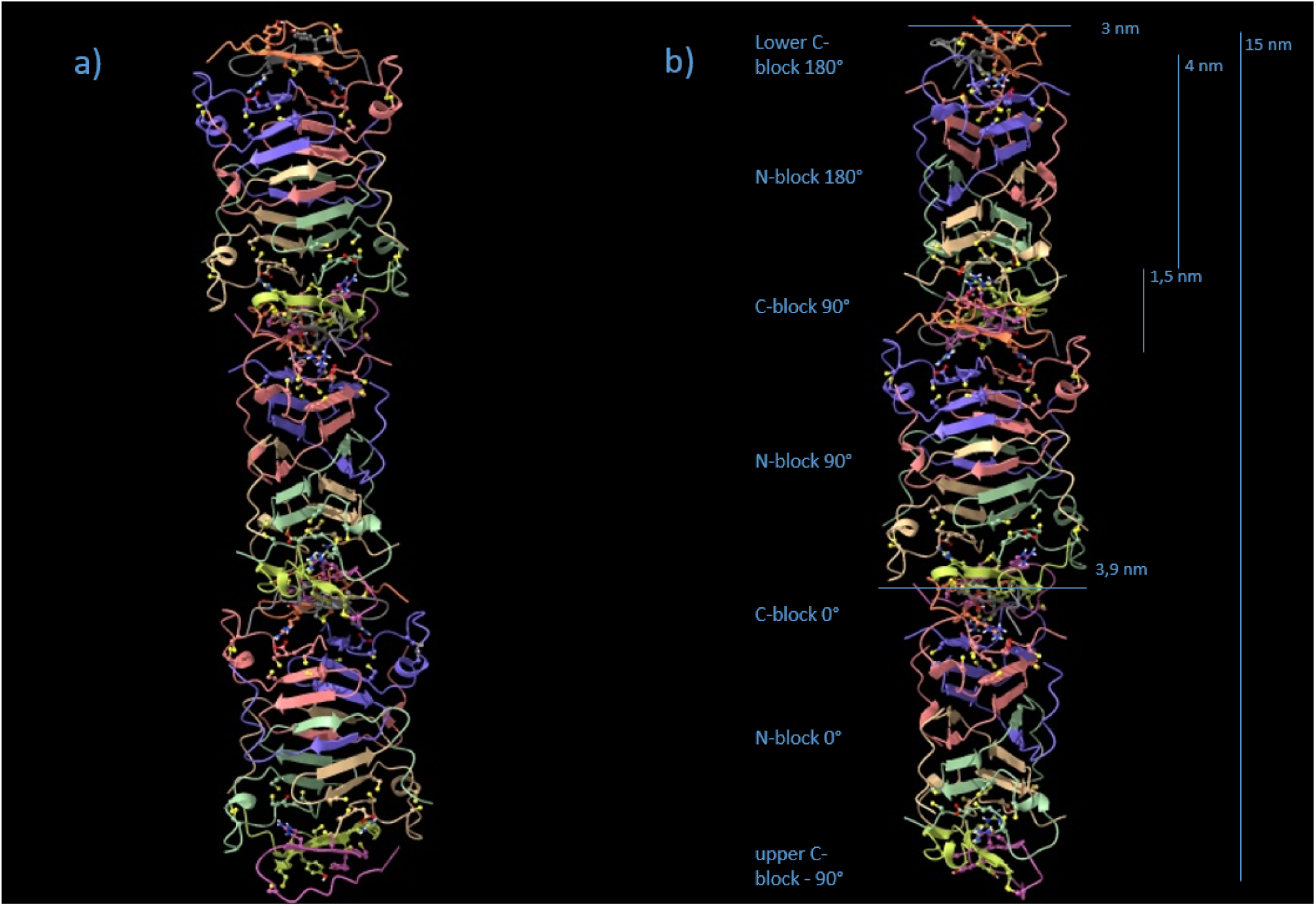
Stacking of F-Keratin into filaments. (a) The repeating unit is 10 nm in height, with a sequence of C-block*_n_*, N-block*_n_*, C90(89°)-block*_n_*_+1_, N90(89°)-block*_n_*_+1_, C180(178°)-block*_n_*_+2_, and N180(178°)-block*_n_*_+2_. Because of the D2 symmetry, C180(178°)*_n_*_+2_ and N180(178°)*_n_*_+2_ are in the same orientation as C-block and N-block, on top of each other. (b) Same sequence, 90° side view. Note, that the model is not energy-minimized.

### 2.7 Solid F-keratin

No diffraction data exist for the N-block/C-block filament on its own. Diffraction samples have been solid F-keratin composites. The meridional reflection (9.4 nm) deviates from the 10 nm as calculated from stacking and docking multiple AlphaFold building blocks (Supplementary Materials 5.6, Data S5). One way to reduce the height in the z-direction would be to introduce, by conformational dynamics, a tilt angle in the central regions of the C-block. GROMACS simulations without the outer aromatic part AA 81–88 allowed the octamer to tilt, with the central *β*-sheets of the C-block as the hinge, by more than 45°(Supplementary Materials 5.5, Movie S1). This tilting motion would be confined by the residues of the C-block. The 81-FGYGFGGLGCF motif (Figure 5d), with its multiple small glycine residues, would still tolerate high orientational flexibility for the large aromatic residues. By intercalation and tilting of the aromatic side chains, due to bending of the flexible gly-rich backbone, the C-block could introduce a height difference of *≈* 0.8 nm between both sides, estimated with a distance of 0.5 nm from C*_α_* to the edge of the aromatic side chain. Over a lateral distance of 2.5 nm, this would result in a tilt angle of arctan(0.8*/*2.5) *≈* 18*^◦^*, used as a setpoint for the GROMACS model in Figure 7. This angle cos(18*^◦^*) = 0.95 would be sufficient to achieve a height reduction of 5%, i.e. 10 nm to 9.5 nm (Figure 8). Such tilted configurations, even if they were obtained only temporarily, allow the dynamic positioning of C-discs 9.5 nm apart from their respective N-block origin. As soon as fixation occurs via the two strong electrostatic bonds in the axial direction to the second next neighboring N-block, the maximum stretch of the Gly/Ser/Ala-rich GSA-string fixates the 9.5 nm distance while squeezing the aromatic residues of the compressed C-discs laterally into the space between neighboring filaments. Tilted N-blocks and 18°C-wedges would, however, not be compatible with the intensity distribution of the 4th layer line signal because of the defined orientation of the “hinge” *β*-sheets in the C-wedges and the perfectly defined interface between N- and C-blocks. The D2 symmetry would be lost, an octamer with irregularly tilted N-blocks would not be sufficient to explain the X-ray pattern. The tilted configuration is more likely a dynamic, amorphous, liquid-crystalline or filamentous precursor for the well-aligned and tightly packed filaments in the solid state (see Supplementary Materials 5.6, Data S6 for the .pdb file).

**Figure 8:**
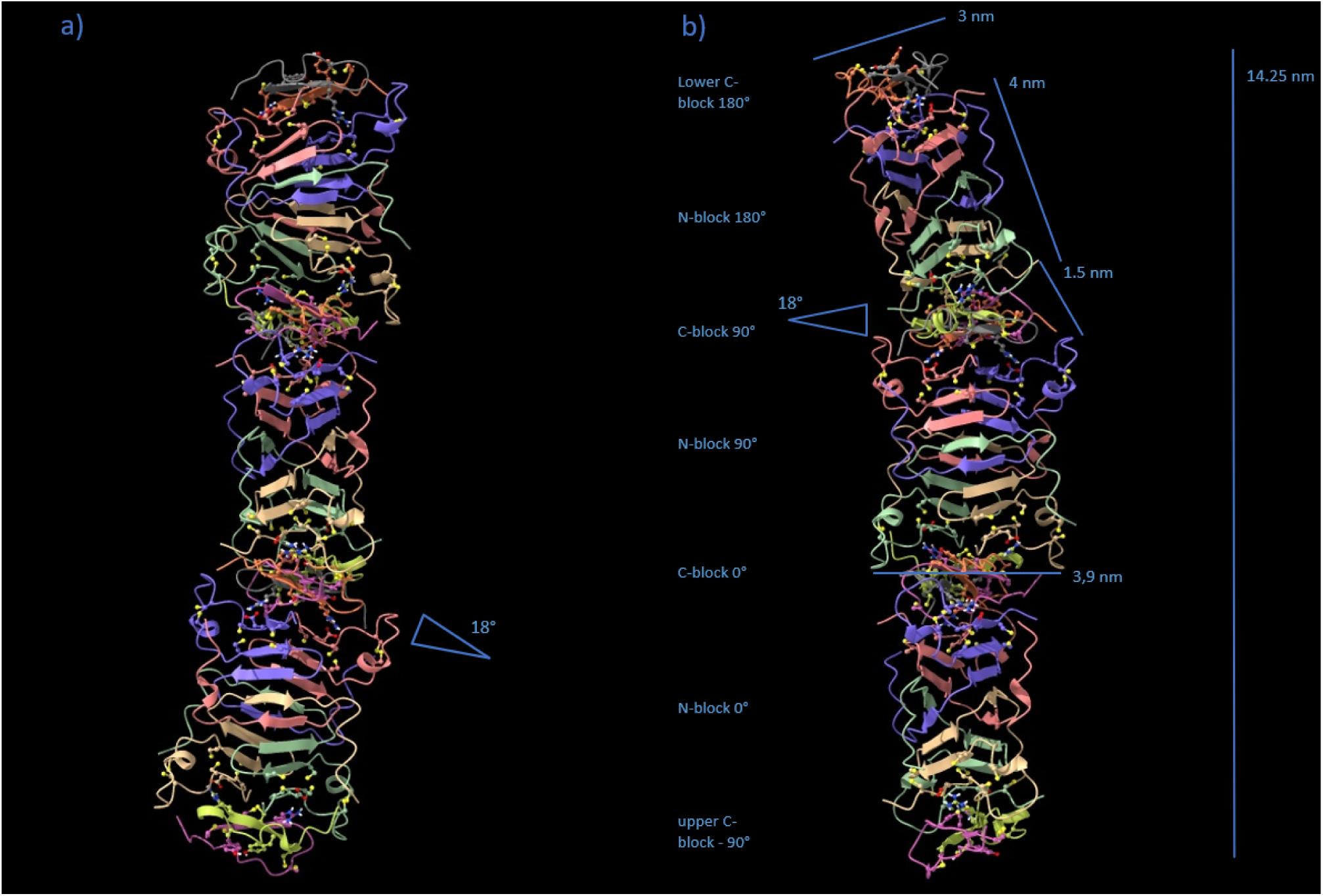
The F-keratin filament in the solid. By temporarily tilting the N-blocks 18°, the straight polymer filament changes to a wiggling geometry. Each two N-blocks touch a slightly squeezed C-block (”C-wedge”) from above and below while turning 90° with respect to each other. The filament curvature allows the most distant axial positioning of C-blocks between N-blocks, with the GSA-strings being fully stretched to 9.5 nm. Note, that the model is not energy-minimized and the GSA-string not shown.

In summary, the obtained results pick up several unique features of solid F-keratin, which no other biological composite material has: The length of the Gly/Ser/Ala-rich GSA-string (”matrix”) is highly conserved, it could be stretched to 9.5 nm, which matches the distance of the repeating unit according to SAXS data (9.4 nm, species-specific). It connects two parts of the F-keratin polymer consisting of 100 amino acids (species specific), the N-terminus and the C-terminus. Folding, according to AlphaFold data, occurs only if 4 units of N-termini and 4 units of C-termini are separately investigated (regardless of species). Tetrameric N-blocks and tetrameric C-blocks reveal dimensions and interfaces that allow them to stably self-aggregate (”polymerize”) into filaments with an axial D2 symmetry and perfect structural fit to SAXS data. C-blocks maintain a certain degree of flexibility so that aromatic side chains would be squeezed out from the main volume of the F-keratin filament, either temporarily or - in the fully solidified 3D arrangement - even permanently. The squeezing of aromatic side chains should affect the lateral arrangements of F-keratin filaments as indicated in the SAXS patterns of solidified peacock rachis.

## 3 Discussion

### 3.1 Previous models of F-keratin

By comparing the complex diffraction pattern of a seagull feather with a simplified pattern after exposing the feather to steam, Fraser et al. [10] concluded in 1971 that the F-keratin is a core-shell composite, in which the outer shell of the cylinder is less stable than the central core, which is built up from pleated *β*-sheets. Up to now, different suggestions were made, such as the *β*-sheets consist either of the AA 22–53 [16] or the AA 27–60 [11, 17–19] segments of the AA 1–100 sequence. In 2010, Pabisch and her colleagues [4] revisited earlier studies [6, 28] and established for the peacock, *Pavo cristatus*, that the axes of a fibrous protein (F-keratin) and the feather rachis are parallel. The radius of the core is remarkably constant over the length of the 1 m long *Pavo* spec. tail feathers (*r_core_* = 0.55 nm) [4]. In contrast, they identified structural changes from the calamus to the central region. The structure is misoriented in the calamus region, but then increasingly aligns and approaches an angle of 90° between axial and lateral reflections. This suggests a tighter packing and would be compatible with a functional contribution of domains (i.e. C-blocks of the new model), which fit increasingly better into the 3D arrangement of N-blocks, which are themselves more tightly packed and better aligned. Likewise, this would also afford the GSA-strings of the new model to co-align. Early X-ray diffraction favored an arrangement of pleated sheets with a helical structure [29, 30]. By a sophisticated analysis of X-ray data, in particular the absence of the first three axial and the visibility of specific odd and even reflections, Fraser et al. [10] developed a model of F-keratin as a pleated sheet framework, where each sheet contains four chains with eight residues per chain and the pairs of sheets are related by a horizontal dyad. To better describe some higher order reflections, an arrangement of the sheets in a right-handed four-fold screw axis was proposed. The length of the repeating unit was 9.46 nm in the seagull feather. Equatorial reflections indicated a layer structure with a lateral period of 3.3 nm. A more detailed model was developed in 2008 [16], with the *β*-sheets densely populated by hydrophobic residues and charged residues, with cysteine residues lying on the outer surface of the filament. In 2021, Parry [12] gives a perfect overview on keratin and intermediate filaments in birds and sauropsids, including a scheme, that shows the arrangement of the *β*-sandwiches with pairs of right-handed twisted antiparallel *β*-sheets, which are related to one another by a perpendicular dyad axis of rotation [12]. This results in a 3.4 nm diameter filament of pitch length 9.6 nm and axial rise 2.4 nm and allows lateral reinforcement of the tissue [12].

### 3.2 The steps forward by the new tentative model of F-keratin

The new tentative F-keratin model is convincing for several reasons. Most importantly, it matches perfectly to the interpretation of the X-ray diffraction data listed in Supplementary Materials 5.4 Table S1.

In the novel AlphaFold-derived model, four chains of either block (N-, C-) are required to form half the repeating unit of the filament. This reminds of the early suggestion of Fraser et al. [10] that the internal arrangement of the filament is a helix with four units per turn of pitch and each *β*-sheet of the pleated sheet framework consists of four chains. Four N-blocks clearly form a core with opposite *β*-sheets, which themselves are rotated according to the AlphaFold-derived model. The novelty here is: This happens as a consequence of four different chains, which intertwine. Most strikingly, the arrangement of these intertwined *β*-sheets matches perfectly to previous data which led to a completely different but still compatible interpretation [12]. Considering only those amino acids which contribute to the *β*-sheet signature of the filament, both models yield a protein density of 1.05 *g/cm*^3^. Despite this apparent similarity between Ref. [16] and the new tentative structure reported here, the origin of each value, however, is fundamentally different (Supplementary Materials 5.6, Table S2). Compatibility with mechanical performance has still to be taken into account (Supplementary Materials 5.2, Appendix A and Appendix B). At present, our model is without alternatives.

### 3.3 The new tentative model: Formation of F-keratin

Polymerization of F-keratin in axial orientation involves ionic fixation between N-block AA 1– 52 and C-block AA 81–100 tetramer interfaces. The external parallelepiped shape of individual N-blocks is reinforced by an unconventional combination of peptide-, disulfide-, and proline-stabilized *β*-sheets which solidify the cross-sectional 3.0 nm *×* 3.9 nm edges and corners of the N/C-tetramer nanoparticles, which form the F-keratin filaments. Additionally, the C-block dimers (= “half-disc”) are covalently fused by disulfide bridges between them, while maintaining a certain degree of shape flexibility using moldable, but sticky aromatic residues. The evolutionary conserved length of the GSA-string AA 53–80 is predestined to enable C-block fusion and disc formation between more distant N-block cores, preventing the structure from disintegrating under strain. The proposed arrangement is a two-phase structure: N/C-tetramers, fixated axially by at least two GSA-strings, provide filament strength. The remaining two strings are free to connect neighboring filaments, if N-block and C-block fold in different filaments. The filament precursor network is then covalently linked via GSA-strings only, without any disulfide cross-linking, forming a palisade-like structure. The low number of linking GSA-strings lead to a weak but clearly visible order in lateral direction, as reflections up to the third order are visible in Figure 3f and 3h, considerably less ordered than in axial direction, where more than 30 layer lines could be identified [10]. In solidified state, the GSA-string (24 vol%; 28% of total AA) which surrounds the N/C-tetramers (76 vol%) is most likely not an amorphous matrix, but stretched and solidified under tension. Forming one stable C-block tetramer may require several intermediate folding steps, accompanied by dynamic N-block tilting and multiple exchanges of C-monomers from different origins. It depends on the availability of C-monomers extending from either the same or two neighboring filaments as a consequence of gradually reaching, in the feather follicle, an intracellular density threshold for the axial packing of either N-blocks or lateral casting of axially aligned filaments. Moldable C-blocks would, step-by-step, occupy the unconstrained 3D arrangement of N-blocks, which get more tightly packed, less tilted and better aligned with time, as soon as most of the C-blocks fall into their respective places. This would agree with the calamus data from the peacock’s rachis as proposed by Pabisch et al. [4].

### 3.4 The new tentative model: Microstructure and mechanics of F-keratin

F-keratin of the peacock feather rachis is a unique material in any respect, there is no other biological material with comparable features. The tentative structure suggests a performance of F-keratin like a metallic super-alloy [31] which is a two-phase single crystal, with a high-volume fraction of hard particles surrounded by a small volume fraction of softer material. Four chains of the N-block intertwine, as well as four chains of the C-block, building up helical arrangements of H-bond stabilized *β*-sheets (Supplementary Materials 5.2, Appendix C). Each pair of blocks is connected by ionic and disulfide bonds in axial orientation and covalently linked to neighboring octamers by four GSA-strings per tetramer unit. This has important mechanical consequences for stabilization and plasticity of the composite [32]. The new tentative model resembles the biomechanics of the ligament-stabilized spinal column of vertebrates, however, on the molecular scale! Densification and biomechanical performance is not taken into account by the AlphaFold approach. This could be a reason for the size deviation of the molecular assembly, derived from AlphaFold (10 nm repeating unit, axial) from X-ray diffraction data (9.4 nm for *Pavo cristatus* [4], 9.6 nm for the seagull feather rachis [9, 12]). Introducing a tilt angle (Figure 8) and preload would not just affect the dimensions of the repeating unit, but introduce further helical features in X-ray diffraction patterns. In general, a mechanical prestress increases the stability, like the spinal ligaments stabilize the spinal column [33, 34]. In F-keratin, this would lead to a reduction of the tetramer size, accompanied by a lateral strutting of aromates, which then interconnect the filaments. One could argue, that not only the absolute precise value of the repeating unit’s size is the criterion to support the model, but much more its specific structural and functional features.

## 4 Conclusion

The new AlphaFold-derived model of avian F-keratin is the first complete description, how the linear sequence of the F-keratin molecule folds to form a solid material with unique, fascinating properties. The F-keratin N-/C-tetramers are arranged like precipitates in a metallic single crystal superalloy, there is a long-range order way beyond the tetramer size. The periodicity of the aligned tetramers is the cause of well-defined diffraction spots. It not only matches perfectly to the interpretation of the X-ray diffraction data and the thus derived models described in the literature, but it assigns each conserved amino acid a specific function for providing a superior combination of mechanical properties. Alpha-Fold directly uses the amino-acid sequences of four partial chains and predicts their positions after folding, whereas the previous models present essentially schemes and cannot unequivocally identify either, the position of amino acids in space or, which sequence or which functional part of the molecular chain was used. The process of structure formation may require temporary tilting of N-blocks and *≈* 5% shortening of the repeating unit in the filaments by dynamic deformation of C-blocks into C-wedges. This is a stable and therefore agreeable transition state in the semi-solid precursor of avian F-keratin. The final solidification step would require 1) N-blocks and C-blocks to align axially, 2) the GSA-strings to maintain a fully stretched position, at least axially, and 3) the aromatic residues to completely squeeze out from the C-wedges into the lateral space between F-keratin filaments or F-keratin sheets. This model is in sufficient agreement with small-angle X-ray data obtained from avian F-keratin. It also explains the mechanical key features of the peacock’s feather rachis. Calculated densities for the solid N-terminal parallelepiped, the flexible C-terminal aromatic sandwich, and the unfolded peripheral GSA-strings would well agree with previously reported experimental data. The *≈* 180° (178°) rotation of 20 intertwisted *β*-strand levels (N-blocks: 16 levels, C-block: 4 levels; *≈* 90° (89°) turn per homo-tetrameric N-core) over 9.4 nm corresponds to an obligate homo-octameric repeating unit.

## Acknowledgements

We thank Josef Lohr who generously provided us with the peacock feather samples. We further thank Silvia Pabisch for her valuable support in X-ray measurements. I.M.W. would like to thank Anna-Katharina Hildebrandt (Dehof) and Andreas Hildebrandt for valuable discussions. The authors gratefully acknowledge the core facility SRF AMICA (Stuttgart Research Focus Advanced Materials Innovation and Characterization) at the University of Stuttgart. We would like to express our gratitude to the communities behind the multiple open-source software packages that were used in the creation of this work.

## 5 Supplementary Materials

Supplementary files are available for download from the Data Repository of the University of Stuttgart (DaRus) at https://doi.org/10.18419/darus-4451

### 5.1 Materials and methods

The F-keratin sequences of the peacock (*Pavo cristatus*), chicken (*Gallus gallus*), and silver gull (*Larus novaehollandiae*) were analyzed as listed in the UniProt database www.uniprot.org under the accession numbers `A0A8C9L9X5_PAVCR`, `P02450_KRFC_CHICK`, and `P02451_ KRFT_CHRNO`. Preliminary 3D predictions for chicken and silver gull can be found in the AlphaFold database www.alphafold.ebi.ac.uk/entry/P02450 www.alphafold.ebi. ac.uk/entry/P02451 (date of access: 29.04.2024).

Sequence alignments were performed using www.uniprot.org/align and www.ebi.ac.uk/jdispatcher/msa/clustalo. Structures were aligned with www.rcsb.org/alignment with jFATCAT (rigid).

Arbitrarily modified F-keratin sequences were used to analyze comparatively any possible effects of point mutations. Sequences were parsed into segments by systematically varying initial points and length. Segments were selected by suspected amino-acid functionality as well as random segments for reference. Sequence segments were analyzed as monomers, dimers, trimers, tetramers, and selected multimers by using Colab-fold (Alpha Fold 2, multimer V3 set) [35].The MSA mode was set to MMseqs2, with par mode set to unpaired+paired in five cycle steps. The obtained structural models were energy minimized to obtain optimized Ramachandran angles, selected according to pLDDT score and distributions, as well as PAE plots. Further judgement criteria were size, shape, density, positioning of functional amino acids, and interfacial configurations of the structural models. Three species were comparatively investigated unless similar results were obtained. Due to various combinations, several hundred input sets were created for carrying out the AlphaFold analyses. Only the best options that led to promising results were carried on. Careful inspections of intermediate results for chemical and functional distinction were mandatory.

The ClusPro server at www.cluspro.bu.edu/home.php [36–39] was used for docking experiments to analyze structural models of selected segments. Initial arrangements were defined according to the best-fit with X-ray diffraction data in lateral and axial orientation. The ClusPro server generally returned 90 to 120 different docking results, with different force modes (balanced, hydrophobic, electrostatic, and van der Waals+electrostatic). A search for structures that would be eventually compatible with experimental X-ray data yielded, all in all, about 4000 results.

The stability of structural models, derived with and without docking, was analyzed with the help of GROMACS 2024 (available at www.gromacs.org) [40–43] and the charm36 (jul2022) force field from www.charmm.org [44–47]. Simulations were carried out using the md-integrator in neutralized systems. Sample MD settings are included (Supplementary Materials 5.6, data S7, S8, and S9)).

ChimeraX, available at www.cgl.ucsf.edu/chimerax/download [48–50], was applied to visualize, measure, and build .pdb-files of the structural models. Specifically, it was used to view our simulations and their preparations and to predict and display structural features, biochemical properties, and molecular interactions (hydrophobic, electrostatic, H-bonds, etc.). Complementary for visualization and modeling the VMD software, available at www.ks.uiuc.edu/Research/vmd [51], was used.

Ramachandran angles and plots were calculated with the Ramachandran-Plotter python code, available at https://github.com/Joseph-Ellaway/Ramachandran_Plotter.

### 5.2 Supplementary text

#### Appendix A: Mechanical data

The excellent mechanical properties of F-Keratin must be the consequence of its nanostructure, its arrangement of peptide bonds, and specific amino acid interactions. Apart from peptide bonds (2.0-4.6 eV), disulfide bonds (2.2 eV) are the strongest bonds in proteins, which often cross-link multiple polymer chains. Other bonds, eg. electrostatic (0.32 eV), and H-bonds (0.1-0.4 eV) are far weaker [21–23, 52–59]. F-keratin contains 6 cysteines within the first 30 amino acids. Hypothetically, three disulfide bonds would amount to 6.6 eV, suggesting that the N-block has the highest bond energy concentration in the whole protein. Any convincing model has to agree with the extraordinary strength of feathers with an activation enthalpy of H = 1.75 eV in an activation volume of v = 0.83 nm^3^ [32, 60, 61], which implies the stiffness of the keratin molecule itself. It was overlooked that cysteines might be part of the *β*-sheets, which were considered to be of primordial importance [10–19].

#### Appendix B: Structural analysis of calculated AlphaFold model

In terms of structural properties, one has to take into consideration that not only the strength of bonds but also the flexibility and dynamics of the polypeptide chain play a major role. Due to the partial *π*-character of the peptide bond, the distribution of *ϕ*- and *ψ*-angles is characteristic of proteins with high *β*-sheet content [62]. Furthermore, prolines could enhance the stiffness of proteins by reducing the degree of rotational freedom of peptide bonds. The N-block of F-keratin contains 8 Pro residues. Their function remains a matter of speculation until today. Ramachandran angles are listed in the Supplementary Materials 5.4, Table S3 and Table S4. Graphical Ramachandran plots are displayed in the Supplementary Materials 5.3, Figure S1 and Figure S2.

#### Appendix C: Functional features of the tentative F-keratin structure

The stiff structural part of the avian F-Keratin protein is a symmetric arrangement of stacked helical *β*-sheets that originate from the intercalating N-termini of 4 individual monomers. Each monomer contributes six cysteines which stabilize the tightly packed secondary structures by disulfide bonds. This arrangement enforces the formation of a proline-stabilized wing screw-like formation, extending at the corners of the tetrameric building block. Hydrophobic residues from all 4 chains are located in the axial core of the parallelepiped-shaped unit, which is further stabilized by one central crossing-over of the 4 polypeptides’ “backbones”. The tetramer of the C-termini forms a sandwich-like disc that almost perfectly fits the parallelogram-shaped (”X-type”) cross-section of the N-terminal tetrameric parallelepiped in size and shape. Aromatic Phe and Tyr residues, contributed from two dimers on each side, are accumulated in the center of the disc. The upper and lower interfaces of the C-terminal tetramer match perfectly with the upper and lower interfaces of the N-terminal tetrameric parallelepiped, stabilized by ionic bonds between the 5-D and 95-R of heterodimers. Tight stacking of two tetrameric N-terminal parallelepiped + C-terminal disc units enables the formation of a filament with 10 nm repeating units and a rhombohedral cross-section with 3.0 nm and 3.9 nm edge length. The directionality of the tetrameric unit in terms of the left-handed helicity of 8 *β*-strands (11.25°per turn) results in a *≈* 90°(89°) turn per tetramer or, *≈* 180° (178°) turn / 10 nm pitch. Another characteristic feature explains the remarkable mechanical strength under tensile load: The 4 unfolded parts (AA 53–80) termed GSA-strings which covalently connect the tightly folded N- and C-terminal tetramers would, if fully stretched, be sufficiently long to span the 9.5 nm axial distance between two repeating units per octamer. The lateral exchange of C-discs between F-keratin filaments holds them in place and horizontally aligns the filaments in sheet-like configurations.

### 5.3 Supplementary figures

Figure S1: N-block Ramachandran (Φ, Ψ) plot of *Pavo cristatus* F-keratin

Figure S2: C-block Ramachandran (Φ, Ψ) plot of *Pavo cristatus* F-keratin

### 5.4 Supplementary tables

**Table S1:** X-ray data and features of the new model.

| Axis | distance | assignment | new model, this publication |
| --- | --- | --- | --- |
| axial | 9.4 nm | F-keratin filament, periodicity of repeating unit | uncertainty in measurements: 9.8 nm - 10.4 nm, in a relaxed state; 9.4 nm - 9.6 nm by tilting |
| lateral | 3.4 nm | distances between core filament axes | lateral distance: 3.4 nm (staggered configuration) |
| axial | 2.35 nm | F-keratin filament, 4th layer line | original dimers: 2.5 nm; height of half a N-block tetramer with half a C-disc |
| lateral | 1.1 nm | N-block unit: inter-sheet distance | stretched GSA-string; or: distance between paired β-strands in the 12 N-block and/or C-block levels |
| axial | 90°/unit | offset, F-keratin core filament | combination of rotation and offset: $\approx 180^\circ(178^\circ)/\text{unit}$, with units twice as large (octamer); (D2 symmetry) |

**Table S2:**
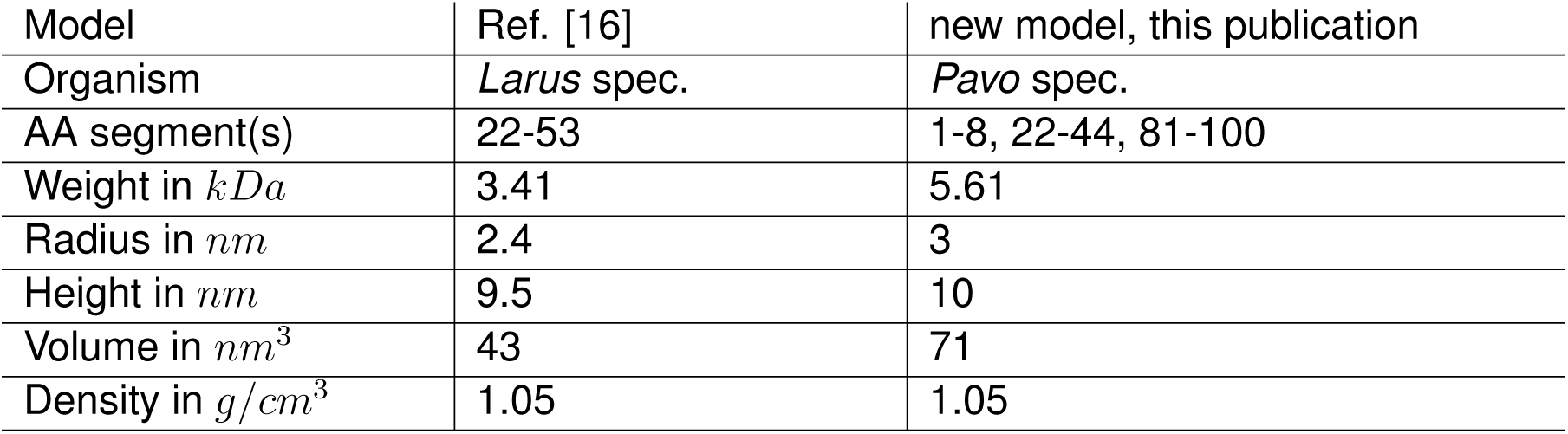
Density calculations.

Table S3: N-block Ramachandran (Φ, Ψ) angles of *Pavo cristatus* F-keratin.

Table S4: C-block Ramachandran (Φ, Ψ) angles of *Pavo cristatus* F-keratin

### 5.5 Supplementary movies

Movie S1: Tilting potential of the fiber model with the N-block AA 1–53 and C-block AA 89–100.

### 5.6 Supplementary data

Data S1: *Pavo cristatus* N-block pdb-file

Data S2: *Gallus gallus* N-block pdb-file

Data S3: *Larus novaehollandiae* N-block pdb-file

Data S4: *Pavo cristatus* C-block pdb-file

Data S5: *Pavo cristatus* F-keratin filament pdb-file

Data S6: *Pavo cristatus* F-keratin filament precursor pdb-file

Data S7: Settings file for ionic equilibration mdp-file

Data S8: Settigs file for md equilibration mdp-file

Data S9: Settings file for md runs mdp-file

## References

1. Terrill RS, Shultz AJ. Feather function and the evolution of birds. Biol Rev Camb Philos Soc 2023, 98:540–66. DOI: 10.1111/brv.12918.

2. Chuong CM, Chodankar R, Widelitz RB, Jiang TX. Evo-devo of feathers and scales: building complex epithelial appendages. Curr Opin Genet Dev 2000, 10:449–56. DOI: 10.1016/S0959-437X(00)00111-8.

3. Greenwold MJ, Sawyer RH. Linking the molecular evolution of avian beta (β) keratins to the evolution of feathers. J Exp Zool B Mol Dev Evol 2011, 316:609–16. DOI: 10.1002/jez.b.21436.

4. Pabisch S, Puchegger S, Kirchner HOK, et al. Keratin homogeneity in the tail feathers of Pavo cristatus and Pavo cristatus mut. alba. J Struct Biol 2010, 172:270–5. DOI: 10.1016/j.jsb.2010.07.003.

5. Busson B, Engström P, Doucet J. Existence of various structural zones in keratinous tissues revealed by X-ray microdiffraction. J Synchrotron Radiat 1999, 6:1021–30. DOI: 10.1107/S0909049599004537.

6. Astbury WT, Marwick TC. X-ray interpretation of the molecular structure of feather keratin. Nature 1932, 130:309–10. DOI: 10.1038/130309b0.

7. Bear RS. Long X-ray diffraction spacings of the keratins. J Am Chem Soc 1943, 65:1784–5. DOI: 10.1021/ja01249a509.

8. Bear RS. X-ray diffraction studies on protein fibers. II. Feather rachis, porcupine quill tip and clam muscle. J Am Chem Soc 1944, 66:2043–50. DOI: 10.1021/ja01240a013.

9. Bear RS, Rugo HJ. The results of x-ray diffraction studies on keratin fibers. Ann N Y Acad Sci 1951, 53:627–48. DOI: 10.1111/j.1749-6632.1951.tb31964.x.

10. Fraser RD, MacRae TP, Parry DA, Suzuki E. The structure of feather keratin. Polymer 1971, 12:35–56. DOI: 10.1016/0032-3861(71)90011-5.

11. Parry DAD. Structure and topology of the linkers in the conserved lepidosaur β-keratin chain with four 34-residue repeats support an interfilament role for the central linker. J Struct Biol 2020, 212:107599. DOI: 10.1016/j.jsb.2020.107599.

12. Parry DAD. Structures of the ß-keratin filaments and keratin intermediate filaments in the epidermal appendages of birds and reptiles (sauropsids). Genes 2021, 12. DOI: 10.3390/genes12040591.

13. Fraser RDB, Parry DAD. Filamentous structure of hard *β*-keratins in the epidermal appendages of birds and reptiles. In: Fibrous Proteins: Structures and Mechanisms. Ed. by Parry DAD, Squire JM. Springer, 2017:231–52. ISBN: 978-3-319-49674-0. DOI: 10.1007/978-3-319-49674-0_8.

14. Fraser RD, MacRae TP, Suzuki E. Structure of the alpha-keratin microfibril. J Mol Biol 1976, 108:435–52. DOI: 10.1016/s0022-2836(76)80129-5.

15. Fraser RD, Parry DA. The molecular structure of reptilian keratin. Int J Biol Macromol 1996, 19:207–11. DOI: 10.1016/0141-8130(96)01129-4.

16. Fraser RDB, Parry DAD. Molecular packing in the feather keratin filament. J Struct Biol 2008, 162:1–13. DOI: 10.1016/j.jsb.2008.01.011.

17. Fraser RDB, Parry DAD. The structural basis of the filament-matrix texture in the avian/reptilian group of hard β-keratins. J Struct Biol 2011, 173:391–405. DOI: 10.1016/j.jsb.2010. 09.020.

18. Fraser RDB, Parry DAD. The structural basis of the two-dimensional net pattern observed in the X-ray diffraction pattern of avian keratin. J Struct Biol 2011, 176:340–9. DOI: 10.1016/j.jsb.2011.08.010.

19. Fraser RDB, Parry DAD. Lepidosaur ß-keratin chains with four 34-residue repeats: Modelling reveals a potential filament-crosslinking role. J Struct Biol 2020, 209:107413. DOI: 10.1016/j.jsb.2019.107413.

20. Jumper J, Evans R, Pritzel A, et al. Highly accurate protein structure prediction with AlphaFold. Nature 2021, 596:583–9. DOI: 10.1038/s41586-021-03819-2.

21. Burley SK, Petsko GA. Aromatic-aromatic interaction: a mechanism of protein structure stabilization. Science 1985, 229:23–8. DOI: 10.1126/science.3892686.

22. Rahman MM, Muhseen ZT, Junaid M, Zhang H. The aromatic stacking interactions between proteins and their macromolecular ligands. Curr Protein Pept Sci 2015, 16:502–12. DOI: 10.2174/138920371606150702131516.

23. Bootsma AN, Doney AC, Wheeler SE. Predicting the strength of stacking interactions between heterocycles and aromatic amino acid side chains. J Am Chem Soc 2019, 141:11027–35. DOI: 10.1021/jacs.9b00936.

24. Andersen SO. Amino acid composition of spider silks. Comp Biochem Physiol 1970, 35:705–11. DOI: 10.1016/0010-406X(70)90988-6.

25. van Rosmalen M, Krom M, Merkx M. Tuning the flexibility of glycine-serine linkers to allow rational design of multidomain proteins. Biochemistry 2017, 56:6565–74. DOI: 10.1021/acs.biochem.7b00902.

26. Holmes KC, Popp D, Gebhard W, Kabsch W. Atomic model of the actin filament. Nature 1990, 347:44–9. DOI: 10.1038/347044a0.

27. Levy ED, Pereira-Leal JB, Chothia C, Teichmann SA. 3D complex: a structural classification of protein complexes. PLoS Comput Biol 2006, 2:e155. DOI: 10.1371/journal.pcbi.0020155.

28. Astbury WT, Beighton E. Structure of feather keratin. Nature 1961, 191:171–3. DOI: 10.1038/191171b0.

29. Schor R, Krimm S. Studies on the structure of feather keratin: I. X-ray diffraction studies and other experimental data. Biophys J 1961, 1:467–87. DOI: 10.1016/S0006-3495(61)86903-8.

30. Schor R, Krimm S. Studies on the structure of feather keratin: II. A beta-helix model for the structure of feather keratin. Biophys J 1961, 1:489–515. DOI: 10.1016/s0006-3495(61)86904-x.

31. Glas R, Jouiad M, Caron P, et al. Order and mechanical properties of the γ matrix of super-alloys. Acta Mater 1996, 44:4917–26. DOI: 10.1016/S1359-6454(96)00096-1.

32. Weiss IM, Schmitt KP, Kirchner HOK. The peacock’s train (Pavo cristatus and Pavo cristatus mut. alba) II. The molecular parameters of feather keratin plasticity. J Exp Zool A Ecol Integr Physiol 2011, 315:266–73. DOI: 10.1002/jez.671.

33. Widmer J, Cornaz F, Scheibler G, et al. Biomechanical contribution of spinal structures to stability of the lumbar spine-novel biomechanical insights. Spine J 2020, 20:1705–16. DOI: 10.1016/j.spinee.2020.05.541.

34. Anderson B, Shahidi B. The impact of spine pathology on posterior ligamentous complex structure and function. Curr Rev Musculoskelet Med 2023, 16:616–26. DOI: 10.1007/s12178-023-09873-9].

35. Mirdita M, Schütze K, Moriwaki Y, et al. ColabFold: making protein folding accessible to all. Nat Protoc 2022, 19:679–82. DOI: 10.1038/s41592-022-01488-1.

36. Desta IT, Porter KA, Xia B, et al. Performance and its limits in rigid body protein-protein docking. Structure 2020, 28:1071–1081.e3. DOI: 10.1016/j.str.2020.06.006.

37. Kozakov D, Beglov D, Bohnuud T, et al. How good is automated protein docking? Proteins 2013, 81:2159–66. DOI: 10.1002/prot.24403.

38. Kozakov D, Hall DR, Xia B, et al. The ClusPro web server for protein-protein docking. Nat Protoc 2017, 12:255–78. DOI: 10.1038/nprot.2016.169.

39. Vajda S, Yueh C, Beglov D, et al. New additions to the ClusPro server motivated by CAPRI. Proteins 2017, 85:435–44. DOI: 10.1002/prot.25219.

40. Abraham M, Alekseenko A, Basov V, et al. GROMACS 2024.2 Manual. 2024. DOI: 10.5281/zenodo.11148638.

41. Markidis S, Laure E, eds. Solving software challenges for exascale. Lecture Notes in Computer Science. Springer International Publishing, 2015. ISBN: 978-3-319-15975-1. DOI: 10.1007/978-3-319-15976-8.

42. Abraham MJ, Murtola T, Schulz R, et al. GROMACS: High performance molecular simulations through multi-level parallelism from laptops to supercomputers. SoftwareX 2015, 1–2:19–25. DOI: 10.1016/j.softx.2015.06.001.

43. Pronk S, Páll S, Schulz R, et al. GROMACS 4.5: a high-throughput and highly parallel open source molecular simulation toolkit. Bioinformatics 2013, 29:845–54. DOI: 10.1093/bioinformatics/btt055.

44. Best RB, Zhu X, Shim J, et al. Optimization of the additive CHARMM all-atom protein force field targeting improved sampling of the backbone φ, ψ and side-chain χ(1) and χ(2) dihedral angles. J Chem Theory Comput 2012, 8:3257–73. DOI: 10.1021/ct300400x.

45. Vanommeslaeghe K, Hatcher E, Acharya C, et al. CHARMM general force field: A force field for drug-like molecules compatible with the CHARMM all-atom additive biological force fields. J Comput Chem 2010, 31:671–90. DOI: 10.1002/jcc.21367.

46. Vanommeslaeghe K, Raman EP, Mackerell AD. Automation of the CHARMM General Force Field (CGenFF) II: assignment of bonded parameters and partial atomic charges. J Chem Inf Model 2012, 52:3155–68. DOI: 10.1021/ci3003649.

47. Vanommeslaeghe K, Mackerell AD. Automation of the CHARMM General Force Field (CGenFF) I: bond perception and atom typing. J Chem Inf Model 2012, 52:3144–54. DOI: 10.1021/ci300363c.

48. Goddard TD, Huang CC, Meng EC, et al. UCSF ChimeraX: Meeting modern challenges in visualization and analysis. Protein Sci 2018, 27:14–25. DOI: 10.1002/pro.3235.

49. Meng EC, Goddard TD, Pettersen EF, et al. UCSF ChimeraX: Tools for structure building and analysis. Protein Sci 2023, 32:e4792. DOI: 10.1002/pro.4792.

50. Pettersen EF, Goddard TD, Huang CC, et al. UCSF ChimeraX: Structure visualization for researchers, educators, and developers. Protein Sci 2021, 30:70–82. DOI: 10.1002/pro.3943.

51. Humphrey W, Dalke A, Schulten K. VMD: visual molecular dynamics. J Mol Graph 1996, 14:33–8, 27–8. DOI: 10.1016/0263-7855(96)00018-5.

52. Joesten MD, Schaad LJ. Hydrogen bonding. Dekker, 1974. ISBN: 978–0824762117.

53. Hopfinger AJ. Chemical bonds: Hydrogen bonding. Science 1975, 189:544–5. DOI: 10.1126/science.189.4202.544.

54. Israelachvili JN. Intermolecular and surface forces. Academic Press, 2011. ISBN: 978-0-12-391927-4.

55. Steiner T. The hydrogen bond in the solid state. Angew Chem Int Ed Engl 2002, 41:48–76. DOI: 10.1002/1521-3773(20020104)41:1<48::AID-ANIE48>3.0.CO;2-U.

56. Fersht AR. The hydrogen bond in molecular recognition. Trends Biochem Sci 1987, 12:301–4. DOI: 10.1016/0968-0004(87)90146-0.

57. Nelson DL, Cox MM. Lehninger principles of biochemistry. Macmillan education. W.H. Freeman, 2017. ISBN: 9781319108243.

58. Privalov PL, Khechinashvili NN. A thermodynamic approach to the problem of stabilization of globular protein structure: a calorimetric study. J Mol Biol 1974, 86:665–84. DOI: 10.1016/0022-2836(74)90188-0.

59. Schreiber G, Fersht AR. Energetics of protein-protein interactions: analysis of the barnase-barstar interface by single mutations and double mutant cycles. J Mol Biol 1995, 248:478–86. DOI: 10.1016/S0022-2836(95)80064-6.

60. Weiss IM, Kirchner HOK. Plasticity of two structural proteins: alpha-collagen and beta-keratin. J Mech Behav Biomed Mater 2011, 4:733–43. DOI: 10.1016/j.jmbbm.2011.02.008.

61. Weiss IM, Kirchner HOK. The peacock’s train (pavo cristatus and pavo cristatus mut. alba) I. structure, mechanics, and chemistry of the tail feather coverts. J Exp Zool A Comp Exp Biol 2010, 313:690–703. DOI: 10.1002/jez.641.

62. Lovell SC, Davis IW, Arendall WB, et al. Structure validation by C*α* geometry: *ϕ*,*ψ* and C*β* deviation. Proteins 2003, 50:437–50. DOI: 10.1002/prot.10286.

